# Intratumor Heterogeneity Through the Lens of Gene Regulatory Networks

**DOI:** 10.1101/2025.04.01.646625

**Authors:** Narein Rao, Nicolae Sapoval, Luay Nakhleh, Hamim Zafar

## Abstract

Intratumor heterogeneity (ITH), the presence of heterogeneous genomic and transcriptomic populations in tumor and its implications for the cancer therapies has been one of the focal points of computational cancer biology in the past years. While ITH is well characterized at the genomic level, the regulatory wiring connecting clonal genotypes to transcriptional states remains largely unmapped. To understand whether the evidence for ITH at the gene regulatory level can be inferred from scRNA-seq data, we analyzed scRNA-seq datasets spanning triple-negative breast cancer, colorectal cancer, glioblastoma, and a longitudinal stage IV breast cancer cohort, inferring clone-specific gene regulatory networks (GRNs) and developing quantitative tools to compare them: a Wasserstein distance-based measure of regulatory divergence, a *k*^*th*^ order neighborhood similarity framework distinguishing structural from functional rewiring, bootstrap-based edge confidence, and an integrated regulon classification pipeline. Our analyses demonstrate that tumor clones exhibit reproducibly distinct network architectures, and dominant clones show significantly greater regulatory divergence from normal cells than smaller clones, linking regulatory rewiring to clonal fitness. Treatment-induced remodeling exceeds clonal variation, with rewiring exhibiting structural divergence but functional convergence. Regulon analysis further resolved clone-variable, sample-specific, and broadly conserved programs, some of which containing transcription factors whose target genes independently were associated with patient survival.

## 1 Introduction

Cancer is a complex disease characterized by extensive genomic and transcriptomic diversity [10] occurring both across different cancer types and patients, as well as within individual tumors. The latter form of diversity manifested by the presence of multiple genotypically distinct cellular populations within the same tumor tissue is known as intratumor heterogeneity (ITH) and presents a major obstacle in design and delivery of efficient and personalized cancer therapies [52, 12]. A common approach to characterize ITH is based on the clonal cancer evolution hypothesis [61], which involves identifying the clonal subpopulations of tumor cells typically defined by their somatic single-nucleotide variants (SNVs) [89, 18], copy-number aberration (CNA) profiles [13] or a combination of both [73, 11]. However, while the inference of clonal subpopulations from single-cell DNA data (scDNA) [68, 88, 74, 90] and single-cell RNA data (scRNA) [1, 20] has become increasingly common in modern studies of tumor evolution, the assessment of clone-specific alterations in gene regulatory networks (GRNs) remains limited. In many single-cell RNA-seq cancer studies, CNA inference from epithelial or tumor cells is routinely used to distinguish malignant from non-malignant cells, leveraging large-scale transcriptional shifts associated with chromosomal alterations [78, 69]. However, the tumor subclones defined by these inferred copy-number profiles are rarely further characterized in terms of their GRN activity or regulatory programs. A limited number of studies have performed the general characterization of GRNs [33], explored network-based approaches to quantify ITH [66], and developed methods that integrate protein–protein interaction networks with scRNA-seq expression data to infer context-specific regulatory interactions [91]. Other studies have focused on regulatory changes associated with specific cancer hallmark pathways [32, 80] or have compared tumor-derived GRNs with regulatory networks inferred from normal tissues to identify cancer-specific rewiring of transcriptional programs [16]. However, these approaches largely treat tumors as relatively homogeneous systems and do not explicitly account for the clonal structure of tumors revealed by single-cell data. Consequently, the heterogeneity of transcriptional regulatory networks across tumor clones within the same tumor—or their evolution across different temporal or treatment-associated states—remains largely unexplored, limiting our understanding of how regulatory rewiring contributes to clonal diversification and tumor evolution.

To address these, here we test the hypothesis that clone-specific changes in GRNs can be inferred and quantified from scRNA-seq datasets. In particular, we analyzed 32 scRNA-seq datasets spanning colorectal cancer (CRC, *n* = 6), triple-negative breast cancer (TNBC, *n* = 12), Glioblastoma (*n* = 11) and one stage IV breast cancer cohort comprising three longitudinal samples (pre-treatment, post-treatment 1, and post-treatment 2), quantifying the extent of clonal GRN divergence from normal cell subpopulations; the extent of GRN divergence across clones and its association with clonal fitness; therapy-induced GRN divergence and finally the variation in GRN activity and architecture across clones and tumor samples, their association with pathways and established gene meta-programs. Specifically, we used a Wasserstein distance–based framework to quantify regulatory divergence between clones, measuring the minimal redistribution required to transform one clone’s regulatory profile into another. We also employed *k*^th^ order neighborhood similarity analysis to differentiate between structural and functional rewiring, revealing patterns of regulatory convergence and divergence that are not apparent from conventional gene expression analyses. Using these approaches, we sought to answer three key questions: (1) to what extent do tumor clones exhibit distinct regulatory architectures compared to normal cell populations? (2) how does clonal GRN divergence relate to clonal dominance and therapy-induced changes? (3) how does clone-specific regulatory activity shape functional pathways and gene meta-programs across tumors?

Our analyses across cancer types revealed consistent differences in regulatory interactions between tumor clones and normal cells (∼19% mean edge difference between clones and normal), as well as among tumor clones (∼ 12% mean edge difference across clones), with the differential regulatory interactions supported by high bootstrap confidence. Importantly, clonal GRNs deviated significantly from randomized null models, indicating non-random regulatory structure. We further found that the dominant clone exhibited greater divergence from the normal cell population compared to smaller clones, suggesting a potential association between regulatory rewiring and clonal fitness. In a longitudinal scRNA-seq dataset from a stage IV breast cancer patient undergoing treatment, we observed variability in gene activity (measured by betweenness centrality) across both clones and treatment timepoints. Regulatory divergence was more pronounced across treatment timepoints than between clones, with GRN rewiring exhibiting substantial structural changes but relatively limited functional shifts over time. For TNBC and Gliobastoma, we captured the activity of biologically relevant transcription factor–target (TF–TG) interactions (regulons), which displayed both clone- and sample-specific variation, as well as consistent activity across multiple samples. These regulons were further associated with enriched immune or cell state associated pathways and gene programs, enabling characterization of subnetwork-level differences across samples and clones. Notably, variation in TF–TG activity within gene programs was driven either by particular clones or by broader, sample-level effects. By addressing these questions, our study provides a comprehensive framework for linking regulatory network architecture to functional heterogeneity in tumors, uncovering mechanistic insights into tumor evolution and potential therapeutic vulnerabilities.

## 2 Methods

### 2.1 Datasets and preprocessing

We analyzed 32 single-cell RNA-seq (scRNA-seq) cancer datasets spanning multiple tumor types, including 12 TNBC patients [85, 20, 63], 6 CRC patients [41], 11 glioblastoma patients [58, 81], and one stage IV breast cancer cohort comprising three longitudinal samples (pre-treatment, post-treatment 1, and post-treatment 2) [62].

The TNBC, CRC and Glioblastoma datasets, along with inferred copy number profiles (CNPs), tumor/normal cell annotations, and cancer-type–specific gene meta-programs, were obtained from the Curated Cancer Cell Atlas (3CA) [79]. Clonal annotations were inferred using Leiden clustering applied to the existing copy number profiles, as suggested in inferCNV [1]. CNPs for the stage IV breast cancer cohort were inferred by applying inferCNV with settings as described in the original study [62]. Clusters containing fewer than 50 cells were excluded to ensure robustness of downstream analyses.

All datasets were preprocessed using standard workflows in SCANPY [84], including quality control, normalization, and filtering.

### 2.2 Inference of gene regulatory networks

GRNs were inferred for each clone using CellOracle [36], a framework specifically designed for single-cell transcriptomic data that integrates gene expression with prior regulatory information. We initially evaluated three widely used GRN inference methods—CellOracle, GRNBoost2 (as part of the SCENIC pipeline) [5, 53], and Inferelator [75]—all of which have been benchmarked extensively and shown to perform competitively across diverse settings in single-cell and bulk transcriptomic data [67, 75, 76, 60]. All methods were executed using default parameters. For Inferelator, GRNBoost2-derived networks were used as prior information due to the absence of curated priors. Across multiple TNBC and CRC datasets, GRNs inferred using GRNBoost2 and Inferelator exhibited limited overlap between tumor-wide and clone-specific networks (Supplementary Fig. 9), indicating reduced consistency at the clonal level. This observation is consistent with prior bench-marking studies demonstrating that Inferelator performance can vary substantially across model configurations and is particularly sensitive to the availability and quality of prior information. In particular, Inferelator has been shown to exhibit high variability across different model selection strategies (e.g., BBSR, StARS, AMuSR), with performance depending strongly on preprocessing and prior specification [76]. More broadly, GRN inference methods that incorporate prior biological knowledge have been shown to yield more stable and biologically meaningful networks compared to expression-only approaches such as GRNBoost2 [23]. Consistent with these observations, CellOracle’s integration of prior regulatory information and ensemble regression framework provides improved robustness in single-cell settings. Consequently, all downstream analyses were restricted to CellOracle-derived GRNs. GRNs were generated using CellOracles workflow and only edges with p-value *<* 0.05 were considered.

Each GRN is represented as a directed weighted graph *G* = (*V, E, W*), where *V* denotes the set of genes, *E* denotes the set of regulatory interactions, and *W* denotes edge weights reflecting regulatory strength.

### 2.3 Bootstrap and null model analysis

To assess the robustness of inferred regulatory interactions, we performed bootstrap resampling for each clone. Specifically, 100 subsamples were generated without replacement using fractions *f* ∈ {0.75, 0.9, 0.95} of cells within each clone. GRNs were inferred for each subsample using the same pipeline as the full dataset. Edge support was quantified as the proportion of bootstrap replicates in which a given edge was recovered.

To construct a null model, clone labels were randomly permuted 100 times within each dataset, and GRNs were inferred for each randomized partition. This procedure provides a baseline for assessing whether observed GRN differences arise from structured biological variation rather than arbitrary partitioning.

### 2.4 Graph similarity and regulatory divergence measures

To quantify similarity between GRNs, we employed both global and local measures of network similarity.

#### Global similarity

Global similarity between two GRNs *G*_1_ = (*V*_1_, *E*_1_) and *G*_2_ = (*V*_2_, *E*_2_) was computed using Jaccard similarity of edges:

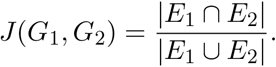

The corresponding Hamming distance was defined as *H* = 1 − *J* .

#### Local similarity

To capture node-level differences, we computed Jaccard similarity over local neighborhoods of genes. For a node *v* ∈ *V*_1_ ∩ *V*_2_, let 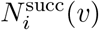 and 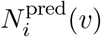 denote its outgoing and incoming neighbors in *G*_*i*_. Local similarities were defined as:

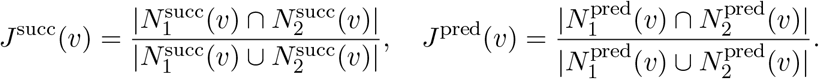

#### Wasserstein distance-based regulatory divergence

To capture graded differences in regulatory strength beyond binary edge overlap, we introduce a weighted divergence measure based on the 1-Wasserstein distance.

For each transcription factor *v* ∈ *V*_1_ ∩ *V*_2_, let **w**_*i*_(*v*) ∈ ℝ*^k^* be the vector of outgoing edge weights from *v* in clone *i*, defined over the union of targets 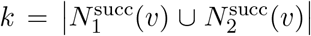, with zero entries for edges absent in a given clone. Let *F*_*i*_(*·*) denote the empirical cumulative distribution function (CDF) of **w**_*i*_(*v*) after ℓ_1_-normalization. The regulatory divergence of TF *v* between the two clones is then defined as the 1-Wasserstein distance between these distributions:

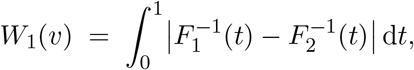

where 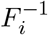 denotes the quantile function of *F*_*i*_. Geometrically, *W*_1_(*v*) is the minimum total “work” required to transform the outgoing regulatory weight profile of TF *v* in clone 1 into that of clone 2. A global summary of weighted regulatory divergence across the network is given by

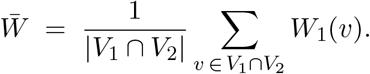

#### *k*-th order neighborhood similarity

For a node *v* and clone *i*, define the *k*-th order successor neighborhood recursively:

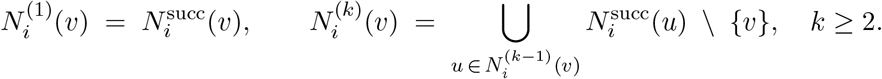

The *k*-th order Jaccard similarity of node *v* between the two clones is then:

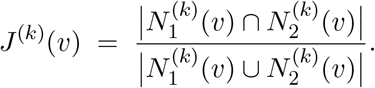

We define the similarity profile of TF *v* up to order *K* as the vector

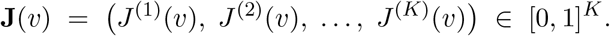

This profile supports a biologically meaningful taxonomy of TF divergence patterns:

1. Structurally divergent, functionally convergent: *J* ^(1)^(*v*) low, *J* ^(*K*)^(*v*) high. The TF drives different direct targets in each clone, but these targets converge on shared downstream effectors.
2. Structurally convergent, functionally divergent: *J* ^(1)^(*v*) high, *J* ^(*K*)^(*v*) low. Direct wiring is similar across clones, but downstream regulatory consequences diverge substantially.
3. Globally divergent: *J* ^(*k*)^(*v*) low for all *k*, indicating pervasive rewiring of the regulatory program of TF *v*.

### 2.5 Centrality analysis

To identify key regulatory genes within the inferred GRNs, we computed degree centrality and betweenness centrality using the NetworkX Python package (v2.8.8) [26].

#### Degree Centrality

The degree centrality of a node *v* ∈ *V* in a network *G* = (*V, E*) is defined as the number of edges adjacent to *v*, normalized by the maximum possible degree:

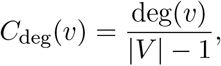

where deg(*v*) is the number of edges adjacent to node *v*. This measure captures the local connectivity of a gene within the regulatory network.

#### Betweenness Centrality

The betweenness centrality of a node *v* ∈ *V* quantifies the fraction of shortest paths between all pairs of nodes that pass through *v*:

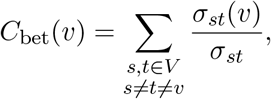

where *σ*_*st*_ is the total number of shortest paths between nodes *s* and *t*, and *σ*_*st*_(*v*) is the number of those paths that pass through node *v*. Betweenness centrality captures the importance of a gene in mediating regulatory interactions across the network.

### 2.6 Regulon activity and pathway enrichment

#### Regulon Activity Scoring

Regulons defined as TF–TG sets derived from either clone-specific GRNs inferred using SCENIC [5] over CellOracle GRN or curated TF–TG interactions from the CollecTRI database [54]. This integration ensures that only biologically supported regulatory interactions are included in each regulon. For each cell in the single-cell RNA-seq dataset, regulon activity scores were computed using the AUCell framework [5] as implemented in the decoupler package (v1.9.2) [6]. Let *G*_*c*_ denote the ranked gene expression vector for cell *c*, and *R* = {*g*_1_, *g*_2_, … , *g*_*m*_} be the set of target genes for a regulon. AUCell computes the area under the recovery curve (AUC) as:

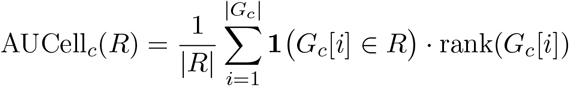

where **1**(*·*) is the indicator function, and rank(*G*_*c*_[*i*]) is the rank of gene *i* in cell *c* (higher rank = higher expression). Higher AUCell scores indicate stronger regulatory activity of the TF in that cell.

#### Regulon Activity Classification

A regulon was considered differentially active across clones if its AUCell scores were significantly different between at least two clones in the same sample (Wilcoxon rank-sum test, *p* < 0.05 after Benjamini-Hochberg correction). Regulons were classified as sample-specific if they showed high activity within a sample but not in other samples. In both clone and sample-specific differentially active regulons, this was enforced by applying restrictions on the number of cells the regulons were active in each group (clone/sample). Specifically, the regulon must be active in at least ℋ % of cells in the group (clone/sample) that the regulon is highly active in and should be active in at most ℒ % of cells in other groups. Depending on the cancer type, ℋ ∈ [40, 50] and ℒ ∈ [20, 35]. Based on these comparisons, three categories of regulons were defined:

- **Clone-specific regulons:** Highly active in a subset of clones within a sample across multiple samples, driving clonal functional heterogeneity.
- **Sample-specific regulons:** Highly active in a sample but not in other samples, reflecting sample-level regulatory programs.
- **Multi-sample regulons:** Observed consistently across multiple samples and clones of a given cancer type, indicating broadly conserved regulatory programs.

#### Functional Enrichment Analysis

To identify biological processes, pathways or meta-programs overrepresented among the genes in each regulon or differentially active regulons, functional enrichment analysis was performed using g:Profiler [39]. Specifically, regulon target genes were tested for enrichment against KEGG [37], HALLMARK [44], WIKIPATHWAY [3]. This approach allowed us to link TF activity patterns to functional programs and interpret the potential phenotypic consequences of clonal regulatory heterogeneity.

### 2.7 Meta-program–specific network analysis

To investigate regulatory structures within biologically relevant gene programs, we leveraged cancer-type–specific meta-programs corresponding to the tumor types analyzed (TNBC and glioblastoma). For TNBC, we utilized predefined meta-programs from the Curated Cancer Cell Atlas (3CA) [79], each comprising approximately 50 genes and associated with malignant cell states, including epithelial–mesenchymal transition (EMT1–3), respiration, translation, cell cycle, stress response, and lineage-specific differentiation amongst others. We first restricted our selection of meta-programs to those prevalent in the TNBC samples considered. We then further filtered the meta-programs to those unique to the samples considered. This left us with 11 meta-programs namely: ‘Cell Cycle - G1/S’, ‘Cell Cycle HMG-rich’, ‘Stress (in vitro)’, ‘Unfolded protein response 1’, ‘Protein maturation’, ‘Translation initiation’, ‘EMT-II’, ‘Interferon/MHC-II (II)’, ‘Epithelial Senescence (HNSCC)’, ‘PDAC-related 5’ and ‘Adherens’. For glioblastoma, meta-programs (39–50 genes each) were obtained from 3CA and the original study by [58], representing developmental cell states such as astrocyte-like (AC-like), oligodendrocyte progenitor-like (OPC-like), neural progenitor-like (NPC-like), and mesenchymal-like (MES-like) programs. Following the same filtering procedure as with TNBC, we were left with 12 meta-programs namely: ‘MES2’, ‘MES1’, ‘AC’, ‘OPC’, ‘NPC1’, ‘NPC2’, ‘G1/S’, ‘G2/M’, ‘Stress (in vitro)’, ‘Interferon/MHC-II (I)’, ‘Interferon/MHC-II (II)’, ‘Glioma single-nucleus’.

For each sample, clone-specific GRNs were inferred using SCENIC in conjunction with CellOracle. For each meta-program, the inferred clone-specific GRNs across all samples were restricted to the subset of genes associated with that program, yielding program-specific subnetworks. Edges within these subnetworks were annotated with bootstrap support values derived from the bootstrap experiment, representing the fraction of bootstrap subsamples in which each edge was observed. This enabled a quantitative assessment of regulatory variability both across clones within a sample and across samples.

To further characterize the role of transcription factors (TFs) within these meta-programs, we evaluated the extent to which they exhibited outlier behavior in degree and betweenness centrality. For each sample, pairwise comparisons were performed across all clone combinations, and a linear model was fit between the normalized centrality values of the corresponding clones. A TF was classified as an outlier if its centrality deviated by at least three standard deviations from the expected value under this model in either clone. An outlier proportion was then computed for each TF as the ratio of the number of clone pairings in which the TF was identified as an outlier to the total number of possible clone pairings within the sample.

This framework enables systematic evaluation of how GRN architecture and transcriptional regulatory activity vary within and across clones and samples, while anchoring these differences to biologically meaningful gene programs.

### 2.8 Cancer Patient Survival Analysis

Patient-level survival analyses were performed using gene expression and clinical data obtained from The Cancer Genome Atlas (TCGA) via the UCSC Xena platform [22]. Normalized gene expression data (RSEM, Toil recompute) and corresponding clinical annotations were retrieved for glioblastoma (GBM) and breast cancer (BRCA) cohorts. Samples were matched across datasets using harmonized TCGA barcodes (first 15 characters), ensuring consistent alignment between expression and clinical records.

For breast cancer, triple-negative breast cancer (TNBC) samples were defined by filtering clinical annotations for estrogen receptor–negative, progesterone receptor–negative, and HER2-negative status. Survival time *T*_*i*_ for each patient *i* was defined as:

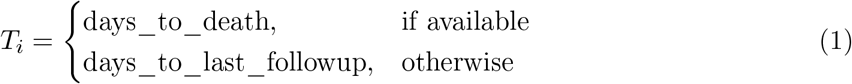

Event status *δ*_*i*_ was defined as a binary indicator of death:

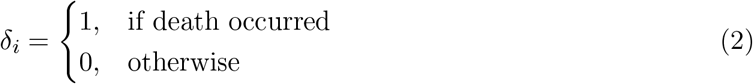

Gene expressions were obtained from matched TCGA bulk samples. Patients were stratified into high and low groups based on gene program enrichment using predefined thresholds (median or quantile-based cutoffs).

Survival distributions for the two groups were estimated using the Kaplan–Meier estimator [38]:

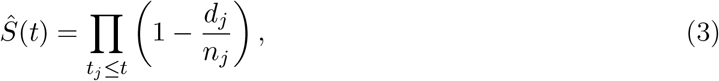

where *d*_*j*_ denotes the number of events and *n*_*j*_ the number of individuals at risk at time *t*_*j*_.

Differences between survival curves were assessed using the log-rank test [48]. For two groups, the test statistic is given by:

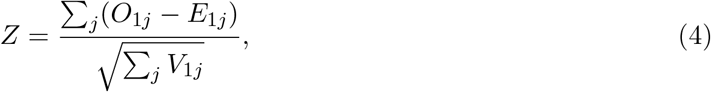

where *O*_1*j*_ and *E*_1*j*_ denote the observed and expected number of events in group 1 at time *t*_*j*_, and *V*_1*j*_ is the corresponding variance. The statistic asymptotically follows a standard normal distribution, from which the corresponding *p*-value was computed. Statistical significance was determined based on the log-rank *p*-value.

## 3 Results

### 3.1 Tumor clones exhibit distinct gene regulatory network architectures compared to normal cell populations

To assess whether tumor clonal GRNs differ from normal cell populations across cancer types, we analyzed 9 TNBC [85, 63] and 6 CRC [41] scRNA-seq datasets, reconstructed the clonal GRNs and compared GRNs inferred from tumor clones and normal cell populations using multiple complementary network similarity metrics. We observed most of the inferred GRN edges to have bootstrap support > 60, allowing us to work with reliable regulatory interactions (Fig. 1d-e). We then compared the edge sets of GRNs inferred from clonal populations and those inferred from normal cell populations (Fig. 1a-c). For each dataset, we evaluated edge-level differences using three complementary strategies: UpSet plots were used to visualize overlaps among GRN edge sets, Jaccard similarity was used to quantify normalized overlap while accounting for differences in network size, and Hamming distance was computed using binarized edge-presence vectors constructed from the union of edges across the compared populations (see Methods). Across all three approaches, clonal GRNs consistently exhibited greater similarity to one another than to networks inferred from normal cells (Fig 1a,b; Supp. Fig. 1a). We calculated the average number of shared edges across clone–clone and clone–normal subpopulation pairs, observing consistently higher overlap among clone–clone subpopulations (Fig. 1a). On average clone-clone GRN difference was 6076 edges (∼ 12%). Furthermore clone–clone pairs showed higher Jaccard similarity than clone–normal pairs across the datasets (Fig. 1b). From our observations we note that clone–clone GRN similarity remained consistently higher than the clone–normal values (Fig. 1b, Supp. Fig. 1a). Hamming distance analysis further supported these differences (Supp. Fig. 1a). Larger clone-clone versus clone-normal differences were associated with disparities in the sizes of the underlying cell populations. For example, samples CID4495, CID4513, CID44971, and TN0131 were outliers among TNBC datasets, while SMC16 and SMC07 were outliers among CRC datasets (Fig. 1a). These samples exhibited substantial differences in the number of cells between normal and clonal populations, without a consistent bias toward either population being larger (Supp. Fig. 2a). Finally, edges unique to clone-specific GRNs tended to have lower bootstrap support compared to the overall set of clone-specific edges (Fig. 1d-e, Supp. Fig. 1b-c). Together, these analyses across both cancer types highlight consistent structural differences between GRNs inferred from tumor clones and those from normal cell populations, supporting the existence of distinct regulatory architectures in tumor clones.

**Figure 1:**
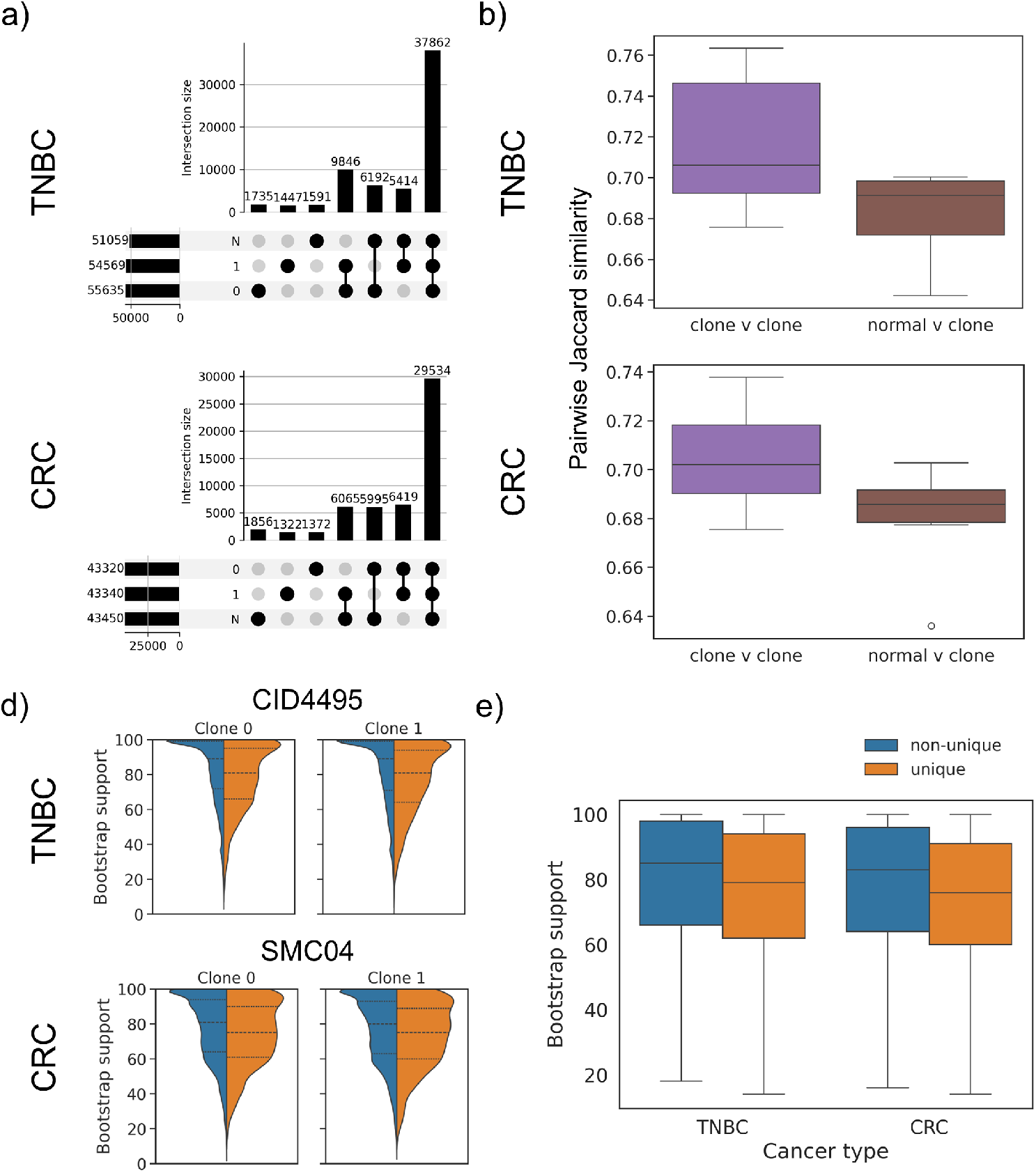
Clonal GRNs exhibit greater structural similarity to each other than to normal GRNs. (a) UpSet plots depicting shared edges between tumor clones and normal cell GRNs for a single TNBC (CID4495) and CRC (SMC04) dataset, highlighting greater edge sharing among tumor clones. (b) Distribution of pairwise Jaccard similarities between edge sets of every pair of clones (purple) and between clones and normal subpopulations (brown). Mann-Whitney U test set to hypothesis with p-values of 0.038 and 0.089 for TNBC and CRC, respectively. (c) Distribution of bootstrap support values for GRN edges not unique to a clone (blue) and edges unique to clone (orange) inferred for TNBC sample CID4495 (top) and CRC sample SMC04 (bottom) for each clone. (d) Distribution of bootstrap support values for GRN edges not unique to a clone (blue) and edges unique to clone (orange) inferred across all TNBC and CRC cancer datasets.

**Figure 2:**
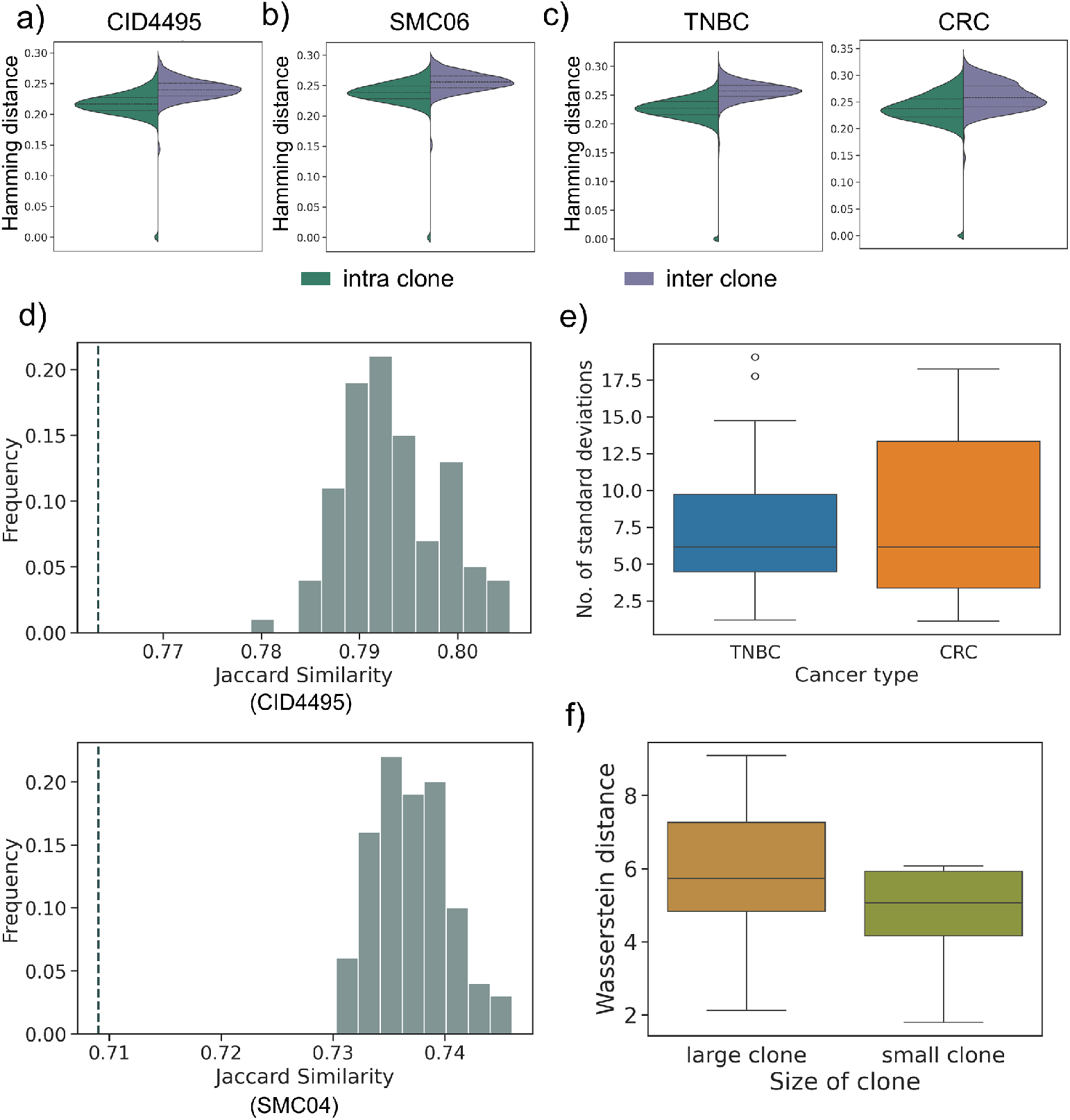
Bootstrap GRNs exhibit stable intra-clone structure, significant inter-clone divergence, with dominant clones exhibiting greater regulatory divergence from normal GRNs. Distribution of Hamming distances between edge vectors of the bootstrap GRNs inferred from sample. Intra-clone (green) distances are measured between all pairs of bootstrap subsamples derived from the same clone, inter-clone (purple) distances are measured between all pairs of bootstrap subsamples derived from different clones. (a) Data for TNBC sample (CID4495) (b) Data for CRC sample (SMC04) (c) Data for all TNBC and CRC samples. (d) Distribution of the Jaccard similarities between edge sets of two tumor subsets obtained by shuffling the cells. Dashed line represent the Jaccard similarity between the edge sets of the clone-specific GRNs. Top: TNBC sample (CID4495). Bottom: CRC sample (SMC04). (e) Distribution of distances between Jaccard similarity of actual clonal GRNs and mean shuffle Jaccard similarity in terms of number of standard deviations across all TNBC and CRC samples. (f) Distribution of Wasserstein distances computed between GRNs inferred from the large clone subpopulation and normal subpopulation (blue); and small clone subpopulations and normal subpopulation (orange) across all 15 datasets (TNBC and CRC). Statistical significance was confirmed using both binomial (*p* = 4.8 × 10^−4^) and Wilcoxon rank tests (*p* = 6.1 × 10^−5^)

### 3.2 Regulatory network divergence distinguishes tumor clones and associates with clonal dominance

In order to establish whether robust GRN differences exists between tumor clones, we compared GRNs inferred from subsampled populations across different clones (inter-clone) with those inferred from subsampled populations within the same clone (intra-clone). To quantify these differences, we computed Hamming distances between binarized edge vectors (edge presence/absence) derived from bootstrap subsamples (subsampling fraction 90%) drawn from the same clones and from different clones (Fig. 2a-c). For both TNBC and CRC datasets, distances between subsamples originating from the same clone were significantly smaller than those between GRNs inferred from subsamples belonging to different clones, indicating consistent regulatory differences between clonal populations. To further confirm that these differences reflect clone-specific regulatory architectures rather than artifacts of sampling variability, we performed a shuffle experiment. Specifically, we compared the similarity between inferred clone-specific GRNs—measured using Jaccard similarity of edge sets—with GRNs inferred from datasets in which clone labels were randomly shuffled within each sample, i.e. a null model (see Methods). Across both cancer types, the Jaccard similarity observed for the original clone-specific GRNs deviated substantially from the distribution obtained from shuffled datasets, with a median deviation of approximately six standard deviations from the shuffled mean (Fig. 2d,e). Although 2 (out of 15) samples (CID44971, SMC16) showed smaller deviations (∼ 2 standard deviations), these corresponded to datasets in which the underlying clones differed substantially in cell counts (e.g., clone1 ≫ clone2) (Supp. Fig. 2a). In such cases, shuffled datasets may still be dominated by cells from one clone, limiting the ability of the shuffle procedure to fully randomize clonal structure. To assess whether GRN divergence is associated with clone size, a proxy for clonal fitness, we quantified regulatory divergence using the 1-Wasserstein distance (Methods) between GRNs inferred for the largest and smallest clonal subpopulations relative to the GRN of the corresponding normal cell population across all 15 datasets. On performing a binomial and Wilcoxon rank test, we observed that the divergence of the GRNs of the largest clonal subpopulations from the normal cell subpopulation were significantly more than the smaller clones (Fig. 2f) We verified that this effect is not driven by differences in absolute cell counts. The sizes of the large clonal, small clonal, and normal subpopulations varied across datasets, ensuring that the observed pattern was consistent across diverse sampling scenarios (Supp. Fig. 2b). These results provide evidence that tumor clones exhibit reproducible and biologically meaningful differences in their gene regulatory architectures.

### 3.3 Treatment-driven divergence of gene regulatory networks across longitudinal tumor samples

For the longitudinal data from the stage 4 breast cancer patient, we focused on GRN differences between the pretreatment tumor sample (Pre) and two post-treatment tumor samples: one originating from the vicinity of central necrotic tissue (Post 1), and another containing viable cancer cells collected near the tumor margins (Post 2) [62]. Out of 23,928 edges inferred for the Pre tumor GRN, only 10.7% (4,765) were shared with Post 1, and 15.07% (8,528) were shared with Post 2. In total, 3,328 edges were common to all three tumor-specific GRNs, while 7,691 edges were shared between Post 1 and Post 2. Due to these large differences in the inferred GRNs, many nodes are unique to their respective GRN while still exhibiting high degree (Supp. Fig. 3). Among these nodes, *ETS1, KLF6*, and *HEY1* showed some of the highest degrees in the Post 1 and Post 2 GRNs but were absent in the Pre GRN. Additionally, these nodes were also strongly supported by subsampling bootstrap values, with average support across Pre:Post 1:Post 2 subsamples of 0 : 81.9 : 79.9, 0 : 78.1 : 83.1, and 0 : 59.9 : 84.9, respectively. In contrast, *ZEB1* and *MEF2A* were detected only in the Pre GRN with bootstrap support of 76.7 : 0 : 0 and 62.9 : 0 : 0, respectively. Given the large discrepancies between the inferred GRNs, we further evaluated nodes with high betweenness centrality to identify potential drivers of key differences between the three samples (Supp. Fig. 4). The transcription factor *EGR1* exhibited among the highest betweenness centrality in both the Pre and Post 2 networks. In contrast, *GLI2* showed high centrality in Post 1 but was absent from the GRNs in both Pre and Post 2. Similarly, *PLAGL1* [2, 29] and *ELF1* displayed high centrality in Post 2 but did not appear as nodes in the GRNs inferred from the Pre and Post 1 tumor samples. Unlike the clonal subpopulations within the same sample, the differences in edges between the inferred GRNs for the three samples were substantial. Although the differences across time-point-specific GRNs were apparent, we also sought to examine how clonal structure influences these centrality measures. To do so, we analyzed differences in degree and betweenness centrality across each clone within each time point. We did not observe large differences in degree centrality, as all clones displayed similar degree distributions. However, we identified notable patterns in betweenness centrality (Fig. 3a). Specifically, the betweenness centrality of several outlier genes—including *FOXF2, GLI2, CREM*, and *EGR1* —appeared to be driven by specific clones to varying degrees at each time point. In some cases, only a small subset of clones contributed substantially to the betweenness property of a gene at a given time point (e.g., *FOXF2*). To investigate this more systematically, we adopted a more robust framework to study the trajectory divergence of GRNs across time points. To quantify GRN divergence, we used our 1-Wasserstein distance formulation and calculated (1) the divergence of GRNs at each time point from the GRN inferred from the normal cell subpopulation, denoted as 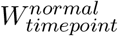 and (2) the divergence between GRNs inferred from different time points, denoted as 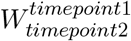 (see Methods). We observed that the Pre time point exhibited the highest divergence from the normal cells, as expected 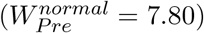, followed by a progressive decrease in divergence from Post 1 to Post 2 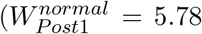, and 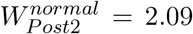 respectively). Next, when examining divergence across time points, we observed higher divergence between Pre–Post 1 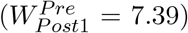 and Pre–Post 2 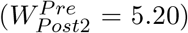 compared to the divergence between Post 1 and Post 2 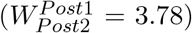. Notably, the magnitude of these time-point divergences (Pre–Post 1 and Pre–Post 2) exceeded the divergence observed among clonal GRNs within the same time point (i.e., GRN divergence between Pre clone GRNs and Post 1 and Post 2 clone GRNs, respectively) (Fig. 3b). Additionally, we also computed the *k*^*th*^-hop jaccard similarity across genes present in the pre and Post 1 time point GRNs; and pre and Post 2 time point GRNs. On calculating the difference between the 0^*th*^-hop jaccard similarity and *k*^*th*^-hop jaccard similarity, we noticed the distribution of differences to incline towards having negative values (Fig. 3c) in both pre vs Post 1 and pre vs Post 2 comparisons. This implies that a large number of interactions across both pre and post time point comparisons showcase structural divergence with functional convergence. Overall, these results suggest that treatment-induced regulatory rewiring may exceed the degree of network rewiring attributable solely to somatic clonal evolution.

**Figure 3:**
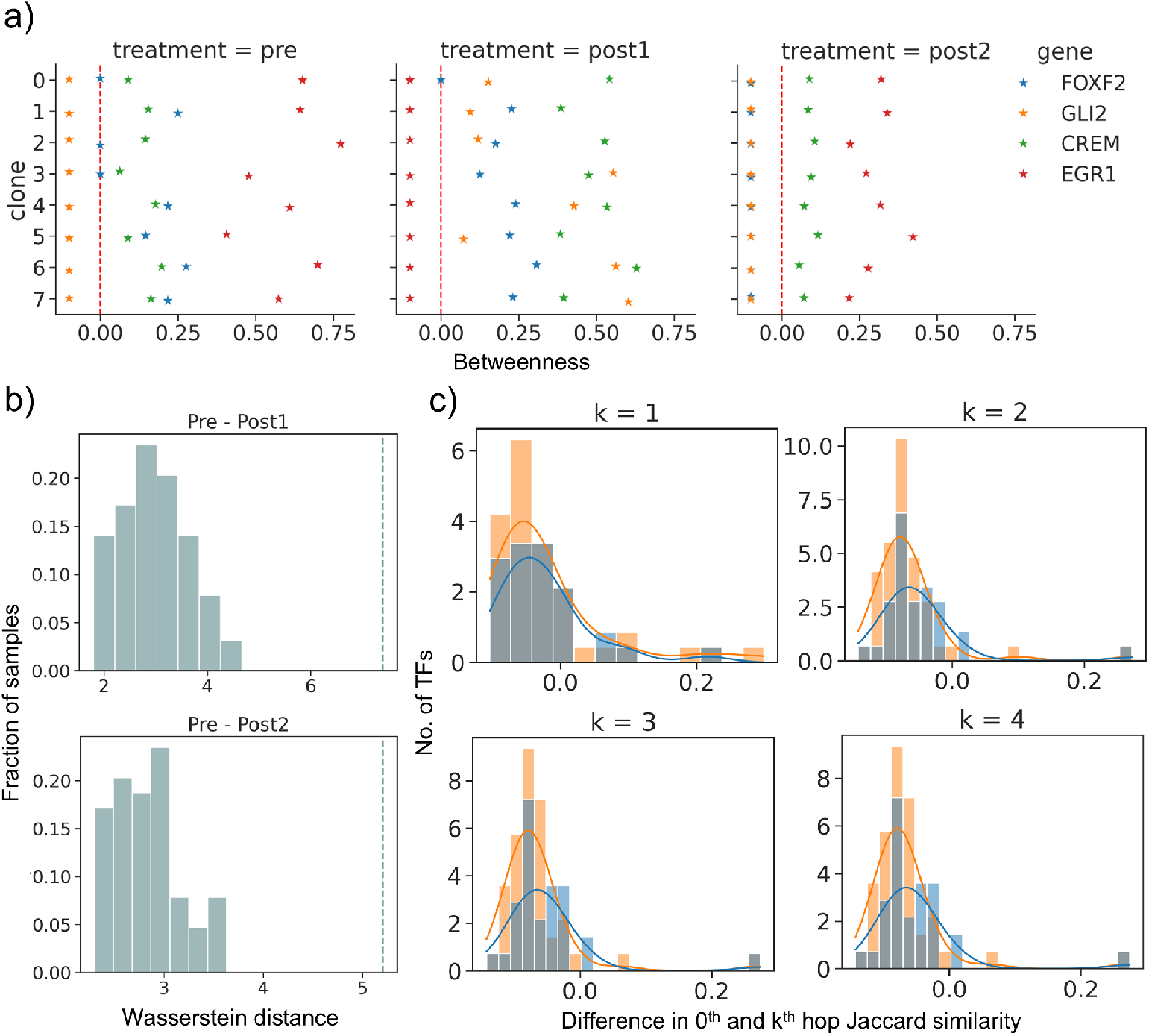
Temporal GRN analysis reveals dynamic, clone-specific centrality shifts, regulatory rewiring across treatment timepoints and structurally divergent network patterns. (a) Betweenness centrality measure for *FOXF2, GLI2, CREM*, and *EGR1* across clones at different time points (pre, Post 1, Post 2). Red dashed line represents 0 betweenness. Genes located to the left of the red dashed line represent isolated nodes within the GRN. (b) Distribution of Wasserstein distance between clone specific GRNs at time points Pre and Post 1 (top); and Pre and Post 2 (bottom). Grey line represents the Wasserstein distance between whole GRNs inferred at time points Pre and Post 1; and Pre and Post 2. (c) Distribution of difference between the 0^*th*^-hop and *k*^*th*^-hop jaccard similarities across common genes in the Pre, Post 1 and Post 2 GRNs. Blue curve represents the comparison between Pre and Post 1 time points and orange curve represents the comparison between Pre and Post 2 time points.

**Figure 4:**
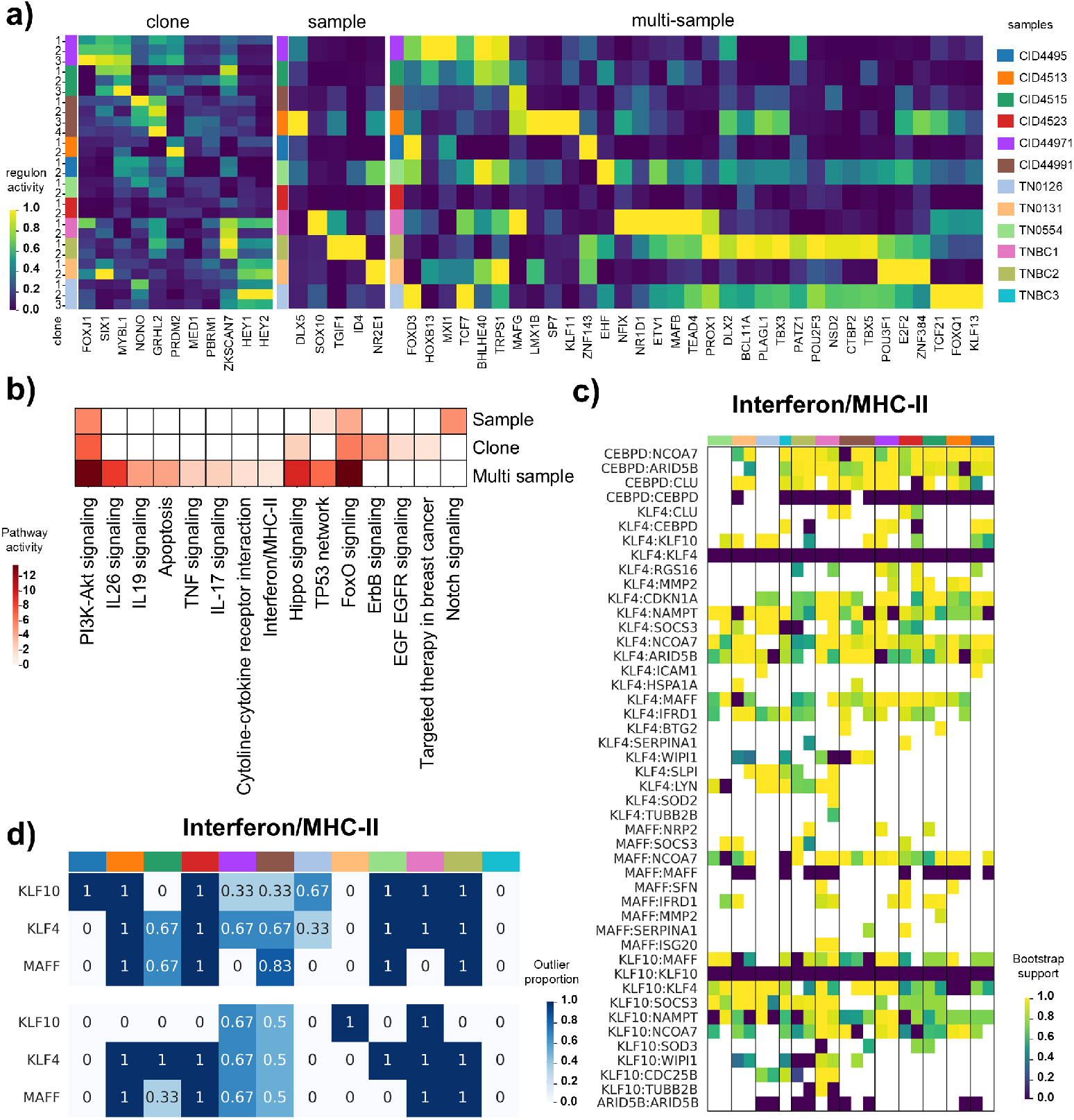
Regulon categories reveal distinct patterns of clone- and sample-specific activity, pathway enrichment, and meta-program–dependent GRN variability across TNBC samples. (a) Activity of three Regulon categories across all TNBC samples: (left to right) regulons that exhibit differential activity across clones in multiple samples, regulons displaying sample-specific activity and regulons that do not vary substantially across clones but show broad activity across multiple samples. x-axis denotes the Regulons, y-axis denotes the samples. (b) Enriched pathways and corresponding pathway enrichment scores for the three regulon categories. Genesets used for pathway enrichment include the TFs and TGs of every regulon within a specific category. (c) Distribution of SCENIC edges across all TNBC samples for meta-program Interferon/MHC-II. Each cell represents the corresponding bootstrap value. Sections within each row (sample) represents clones within each sample. (d) Distribution of degree (top) and betweenness (bottom) outliers, across all TNBC samples for the Interferon/MHC-II meta-program associated TFs. Each cell represents the fraction of clone pairings a gene has been found to be an outlier in.

### 3.4 Regulon activity reveals functional heterogeneity across clones and tumor samples

Regulons, sets of genes regulated by a common transcription factor (TF), play an important role in understanding disease-associated regulatory mechanisms [57, 5, 24]. Since the expression levels of target genes reflect upstream TF activity, regulon analysis provides a useful framework for inferring regulatory activity and characterizing localized GRN dynamics within a given dataset. Having observed clear structural differences across clonal GRNs in multiple cancer types, we next examined regulon activity across clones and samples within each cancer type, followed by pathway enrichment analysis of regulon-associated genes. To ensure reliable regulon definitions, we relied on curated TF–TG interactions from CollecTRI [56]. We first analyzed 12 TNBC samples [85, 63, 20] (see Methods), computed the activity of regulons in each clone of each sample and also identified regulons exhibiting differential activity across clones and samples. Specifically, based on the clonal and sample-level activity, we categorized regulons into three groups: (1) regulons that exhibit differential activity across clones in multiple samples, (2) regulons that do not vary substantially across clones but show broad activity across multiple samples, and (3) regulons showing sample-specific activity (Fig. 4a). Importantly, regulons GRHL2 and SIX1, which are known to be associated with EMT [25, 14] and HEY1 and HEY2, downstream targets of Notch signaling [19] exhibited differential activity across tumor clones in multiple samples. On the other hand, regulons BCL11A and TBX3 (stemness drivers), TRPS1 (marker for TNBC, known to promote epithelial proliferation [4]) and POU2F3 (defines a rare, highly aggressive “Tuft cell-like” subtype of TNBC with a distinct chemosensory gene signature [86]) were found to have activity across all clones in multiple samples.

Notably, some of these regulons contained TFs or TGs that exhibited alterations in copy number profiles (CNP) such as *PBRM1*, suggesting potential links between genomic variation and regulatory activity. To further characterize the functional roles of these regulons, we performed pathway enrichment analysis on the TFs and TGs associated with regulons in each category. While several biologically relevant pathways were shared across samples and clones (e.g. PI3K-Akt signaling pathway [50], FoxO signaling [65]) (Supp. Fig 5a), we also identified pathways uniquely enriched within each regulon category (Fig. 4b) such as EGF EGFR signaling [51] and Notch signaling pathways [21]. These results highlight how distinct sets of regulons may contribute to functional differences across clones or samples. One notable observation was the enrichment of Interferon/MHC-II–related gene meta program within the category of regulons showing broad activity across multiple samples. This raised the question of how GRN structures vary across well-characterized functional meta-programs [79]. To examine the structural impact of clonal variation on these gene meta-programs, we first constructed SCENIC-derived regulatory networks using the inferred CellOracle networks for each clone within every TNBC dataset. We then restricted each network to genes belonging to specific meta-programs associated to the samples considered (see Methods). Fig. 4c illustrates the edges present across clones within each dataset, along with the associated bootstrap support values for each edge for meta-program ‘Interferon/MHC-II’. We observed that some regulatory interactions were present in certain clones but absent in others, or appeared in specific samples but not across the entire cohort. Interactions that varied across clones also included previously known interactions (e.g., *KLF4:SOCS3* [47]). Similarly, we also observed interactions involving *JUND* (a key regulator in TNBC [15, 46]) in the PDAC-related meta-program (Supp. Fig. 6b); and *DDIT3* -mediated interactions in the stress meta-program (Supp. Fig. 6d). Several of these metaprogram-associated TFs exhibited outlier degree or betweenness centrality across samples and clones (Fig. 4d, Supp. Fig. 6). Importantly, some of the TFs (*KLF10* within the Interferon/MHC-II meta-program) or TGs (*PSAT1*, a target of *DDIT3* in the stress-response meta-program) associated with the metaprograms were further found to have significant effects on patient survival in TNBC patients from TCGA cohort [59] (Supp. Fig. 8a).

**Figure 5:**
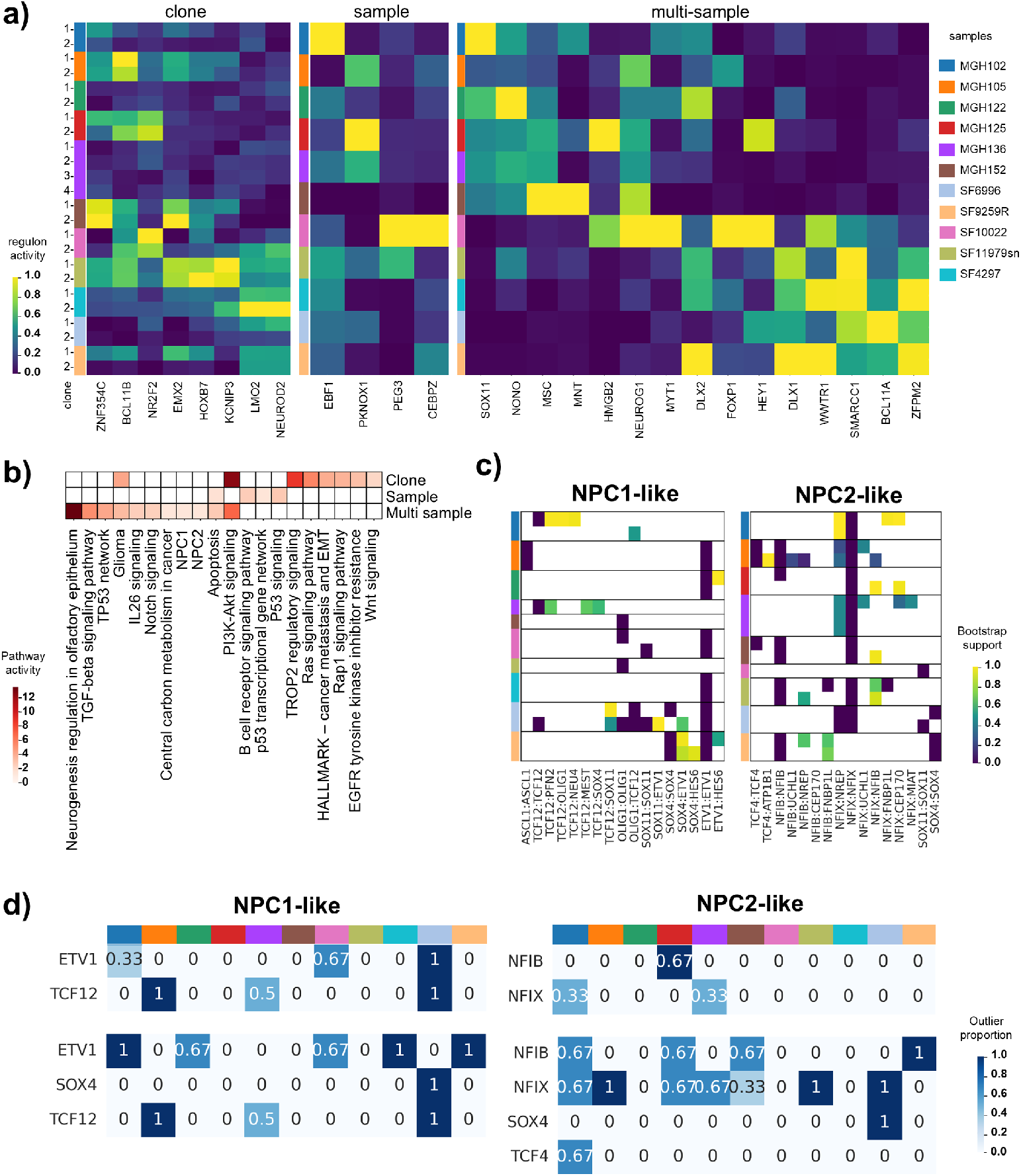
Regulon categories reveal distinct patterns of clone- and sample-specific activity, pathway enrichment, and meta-program–dependent GRN variability across Glioblastoma samples. (a) Activity of three Regulon categories across all Glioblastoma samples: (left to right) regulons that exhibit differential activity across clones in multiple samples, regulons displaying sample-specific activity and regulons that do not vary substantially across clones but show broad activity across multiple samples. x-axis denotes the Regulons, y-axis denotes the samples. (b) Enriched pathways and corresponding pathway enrichment scores for the three regulon categories. (c) Distribution of SCENIC edges across all Glioblastoma samples for meta-programs NPC-1 and NPC-2 respectively. Each cell represents the corresponding bootstrap value. Sections within each row (sample) represents clones within each sample. (d) (left to right) Distribution of degree and betweenness outliers, respectively, across all Glioblastoma samples for NPC-1 TFs and NPC-2 TFs. Each cell represents the fraction of clone pairings a gene has been found to be an outlier in.

### 3.5 Clonal regulatory architecture across glioblastoma meta-programs

We next performed the regulon analysis on 11 glioblastoma samples [58, 81] and again identified three categories of regulons consistent with our observation in TNBC (Fig. 5a). Regulons that showed differential activity across clones in multiple samples included oncogenic drivers HOXB7 (promotes tumor proliferation and associated with Wnt/*β*-catenin pathway) [92], and BCL11B (maintains stemness and helps bypass senescence) [42], as well as the angiogenic factor NR2F2 [70]. Different tumor suppressors such as EBF1 [43] and PEG3 [35] showed sample-specific activities. Regulons which were broadly active across clones and patients included neurodevelopmental TFs (*DLX1, DLX2, SOX11, NEUROG1*) [58], TFs associated with mesenchymal and stemness program (*FOXP1, WWTR1*) [8] and TFs associated with proliferation (*HMGB2*) and chromatin-remodeling (*SMARCC1*) [77]. Interestingly, *SOX11* also exhibited copy-number aberrations in the same datasets where its regulon activity was found to be differentially regulated. Different pathways were enriched for the three classes of regulons (Fig. 5b). TROP2 regulatory signaling [31], Ras signaling [30], Wnt signaling [40] and EGFR tyrosine kinase inhibitor resistance [64] were enriched in the regulons showing differential clonal activity. On the other hand, regulons active across clones and samples enriched for TGF-beta signaling [27], IL26 signaling [87], Notch signaling [7] and Neural Progenitor-like cell state associated pathways [45].

Glioblastoma-related meta-programs [58] were also enriched within the category of multi-sample regulons. In particular, we identified the NPC-1 and NPC-2 meta-programs and examined the structural impact of clonal variation on these gene meta-programs across samples. We observed regulatory edges that were selectively active across clones and samples, with varying bootstrap support values (Fig. 5c). TFs (*TCF12, OLIG1*) involved in these edges are known to play vital roles in gliomagenesis [55]. *TCF12* and *OLIG1* have been reported to be co-expressed within oligodendrocyte progenitor-like lineages in GBM, and *Tcf3* /*Tcf12* double knockout driven by *Olig1* - Cre directly impairs OPC generation and oligodendrocyte differentiation, supporting a functional role for *TCF12* as an E-protein partner within the *OLIG1* -associated transcriptional program [83]. Several of these TFs also exhibited outlier degree or betweenness centrality across clones and samples, suggesting clone-specific regulatory influence within these programs (Fig. 5d). Similar trends were also observed in sub-networks corresponding to other meta-programs in GBM (Supp. Fig. 7). For example, the interaction *DDIT3* :*ATF3* was active in multiple samples for the MES2 meta-program, both factors are known to be co-induced in the pseudopalisading necrotic zones of GBM where ER stress and hypoxia converge to drive the MES-like cellular state [34]. Upon evaluating gene importance in the TCGA GBM dataset, [9] we observed that some interactions involved genes with significant associations with patient survival, including *FTL* (MES-2 meta-program), *SOX11* (NPC-1 and NPC-2 metaprograms) (Supp. Fig. 8b).

## 4 Discussion

In this study, we systematically investigated the impact of ITH on GRN architecture across three cancer types. By inferring clone-specific GRNs from scRNA-seq data across 32 datasets spanning triple-negative breast cancer, colorectal cancer, glioblastoma, and a longitudinal stage IV breast cancer cohort, we demonstrate that tumor clones harbor distinct regulatory architectures that differ from both normal cell populations and from one another, suggesting that regulatory rewiring is an important component of tumor evolution. A central finding of our study is that tumor clones exhibit GRN architectures more similar to each other than to normal cells. These differences were consistent across tumor types, supported by multiple metrics and high bootstrap support, indicating robust and reproducible regulatory interactions. Edges unique to clone-specific GRNs tended to have lower bootstrap support than shared edges, consistent with the expectation that clone-specific regulatory interactions are more labile or context-dependent than widely shared ones.

Another key finding of this study is the association between regulatory divergence and clonal dominance. We developed a novel Wasserstein distance framework that captures graded differences in regulatory strength by comparing distributions of edge weights across transcription factors, thereby providing a global measure of weighted regulatory divergence that complements binary metrics such as Jaccard and Hamming distances. Using this Wasserstein distance framework, we found that the dominant clonal subpopulation within each tumor exhibited significantly greater regulatory divergence from normal cells compared to smaller clones, supported by both binomial and Wilcoxon tests, indicating a mechanistic connection between regulatory rewiring and clonal fitness. We interpret this as clones undergoing greater regulatory reprogramming acquiring transcriptional states that confer selective advantages, potentially through activation of proliferative, anti-apoptotic, or immune-evasive programs [28], consistent with known roles of phenotypic plasticity in cancer [49]. However, regulatory divergence may be either a driver or a consequence of clonal expansion. Longitudinal studies with matched genomic and transcriptomic profiling at the clonal level will be needed to fully disentangle these possibilities [82].

Our analysis of the longitudinal stage IV breast cancer dataset suggests that treatment can induce large-scale regulatory rewiring that exceeds the variation attributable solely to somatic clonal evolution. Several transcription factors with high betweenness centrality emerged as potential drivers of these treatment-associated changes. The clone-specific betweenness analysis further showed that therapeutic pressure acts differentially on the regulatory programs of distinct clonal populations. The k*th* order neighborhood similarity analysis further indicated that many interactions exhibited structural divergence with functional convergence [17] – suggesting adaptive preservation of key transcriptional outputs under therapeutic pressure [49]. However, we acknowledge that inferences from a single longitudinal patient should be interpreted as proof-of-concept, and extension to larger longitudinal cohorts will be essential to establish generalizability.

The regulon-level analyses across TNBC and glioblastoma datasets revealed three distinct categories of regulatory programs: those exhibiting differential activity across clones in multiple samples, those active specifically within individual samples, and those broadly conserved across clones and samples. The pathway enrichment analysis of these regulon categories further revealed that regulatory heterogeneity across clones corresponds to functional differences in tumor biology. The enrichment of immune-related pathways among conserved regulons is particularly notable, as it suggests that the tumor immune microenvironment may exert a consistent selective pressure on regulatory programs across clones and patients, in contrast to the more stochastic activation of specific oncogenic pathways within individual clones [72]. In both TNBC and glioblastoma, regulatory edges within meta-program subnetworks displayed clone- and sample-specific presence with varying bootstrap support, indicating that even within a defined functional program, the regulatory wiring is not static across clonal populations. The identification of transcription factors with outlier degree or betweenness centrality within meta-program subnetworks adds a network-topological layer to functional heterogeneity, highlighting clone-specific regulatory hubs as potential therapeutic vulnerabilities. This complements differential expression analysis by capturing the regulatory logic underlying transcriptional changes. Genes within meta-program subnetworks were associated with patient survival in TCGA cohorts for both TNBC and GBM, indicating that the inferred regulatory interactions reflected biologically and clinically meaningful aspects of disease progression.

A limitation we do acknowledge is that GRN inference from single-cell transcriptomic data remains sensitive to noise and modeling assumptions, and the inferred regulatory interactions will require experimental validation. While our analysis supports regulatory modeling as a feature of tumor evolution, the identification of novel TF-TG interactions is inherently constrained by the capabilities and biases of current state-of-the-art GRN inference methods. In addition, clonal populations were computationally inferred rather than determined through direct lineage tracing, which may introduce uncertainty in clone assignments. Importantly, for a subset of datasets, clonal structure was inferred solely from scRNA-seq data rather than from paired DNA-RNA measurements. This reflects a fundamental limitation: inferring clonal identity from transcriptomic data – the same modality used to derive GRNs – raising a concern of inflated associations between clonal identity and regulatory variation. To mitigate these concerns, we employed multiple safeguards. First, we focused on robust, bootstrap-supported regulatory interactions to reduce sensitivity to stochastic noise. Second, we validated key observations across multiple independent samples and cancer types, ensuring that identified patterns were reproducible rather than dataset-specific artifacts. Third, comparisons against randomized or shuffled controls were used to establish that observed structural differences exceeded those expected by chance. Another limitation is that our analysis focused on transcription factor-target gene interactions and did not account for epigenomic regulation, post-transcriptional mechanisms, or protein-level regulatory layers. Multi-omic approaches integrating single-cell ATAC-seq, proteomics, and spatial transcriptomics with scRNA-seq would enable a more comprehensive characterization of the regulatory landscape underlying intratumor heterogeneity. Spatial transcriptomics in particular could reveal how tumor microenvironment modulates clone-specific regulatory programs [71] – a dimension that is lost in dissociated single-cell assays. Finally, while we analyzed three major cancer types, the generalizability of our findings to other tumor types and histological subtypes remains to be established, and future studies should extend this framework to additional cancers.

In summary, our work provides a comprehensive framework for characterizing intratumor heterogeneity at the level of gene regulatory networks. As single-cell multi-omic technologies continue to mature and computational methods for GRN inference improve, the integration of regulatory network analysis into the standard toolkit for studying intratumor heterogeneity promises to yield deeper insights into the mechanistic basis of tumor evolution and, ultimately, to inform the design of more effective and precisely targeted therapeutic strategies.

## 4.1 Acknowledgements

This research was supported in part by the National Science Foundation, grants IIS-1812822 and IIS-2106837 (L.N.) and the DBT/Wellcome Trust India Alliance grant IA/E/21/1/506298 (H.Z.).

## 4.2 Author Contributions

H.Z. and L.N. conceived the study. N.R., N.S., H.Z., and L.N. developed the methodology. N.R. implemented the software, performed the experiments and ran baseline methods under the supervision of N.S., H.Z., and L.N. N.R., N.S., and H.Z. analyzed the results. N.R., N.S., and H.Z. drafted the manuscript. All authors reviewed, edited, and approved the final version of the manuscript.

