## Supplemental figures for "Intratumor Heterogeneity Through the Lens of Gene Regulatory Networks"

### A Supplementary figures

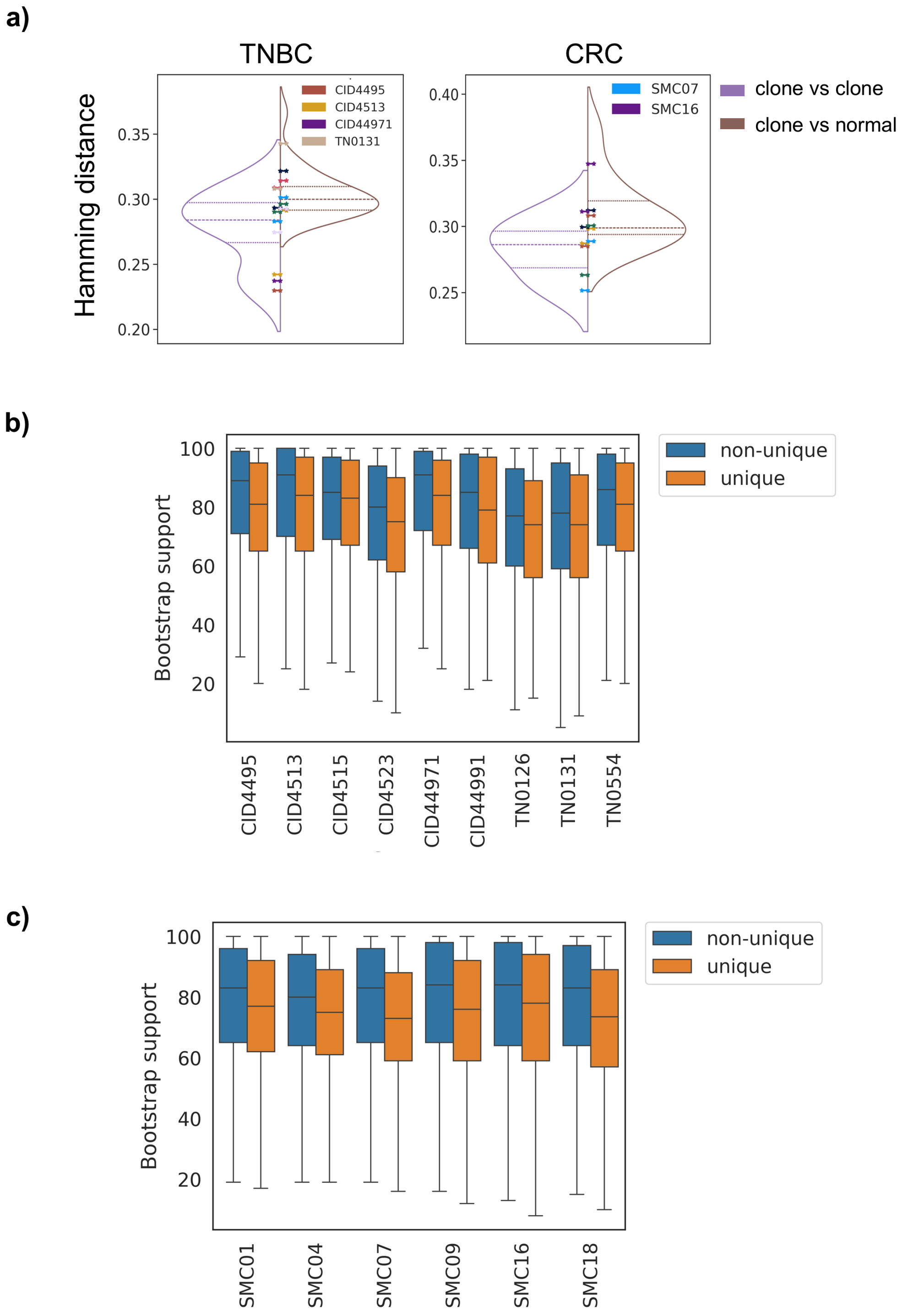


Supplementary Figure 1: (a) Distribution of Hamming distances using binarized edge-presence vectors between clones (purple) and clone and normal subpopulations (brown). Each color peg within the violin plot depicts a specific sample. Mann-Whitney U test set to hypothesis with p-values of 0.0039 and 0.083 for TNBC and CRC, respectively. (b)

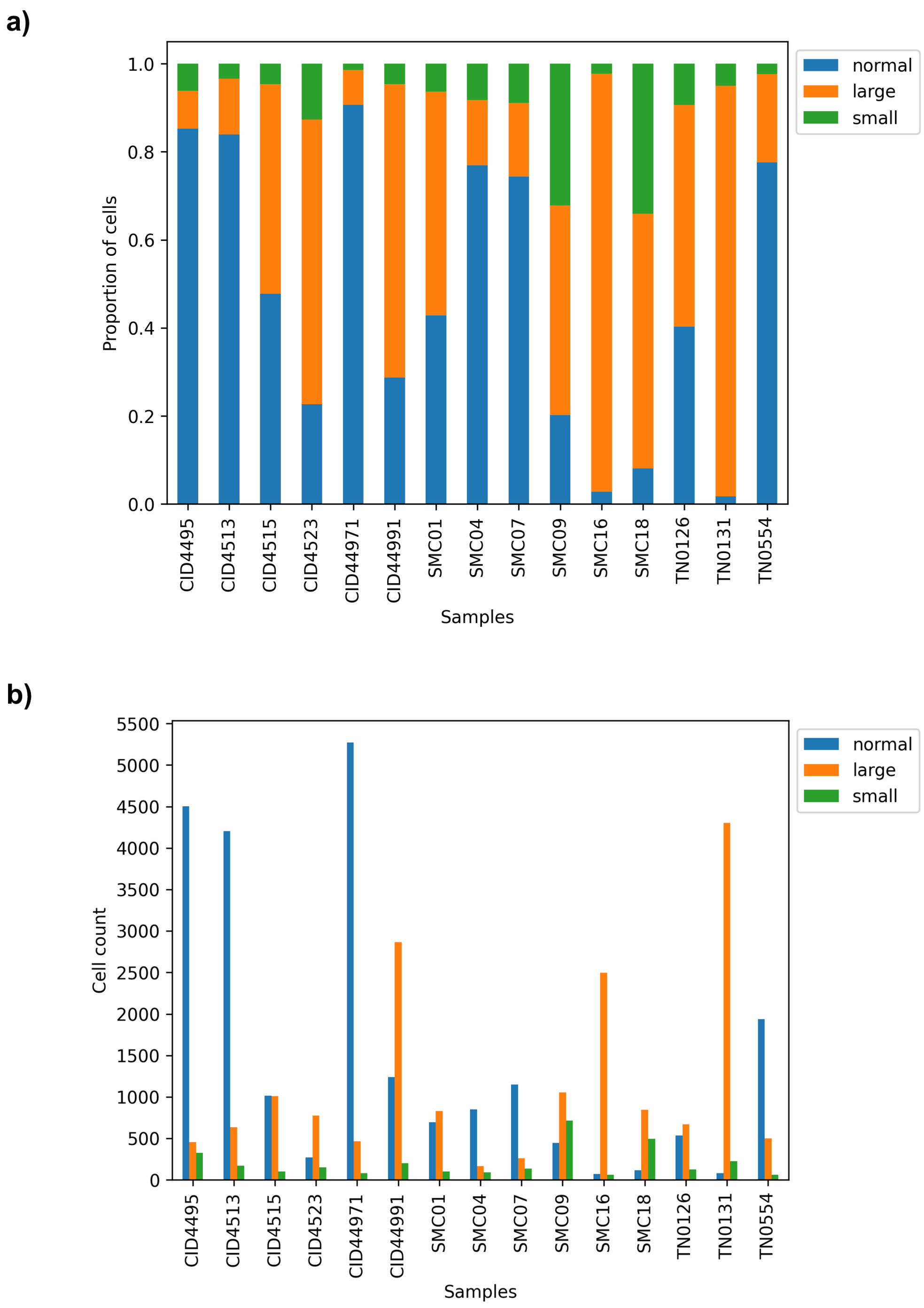


Supplementary Figure 2: (a) Proportion of cells across normal cells, largest and smallest clones in each sample. (b) Absolute counts of cells across normal cells, largest and smallest clones in each sample

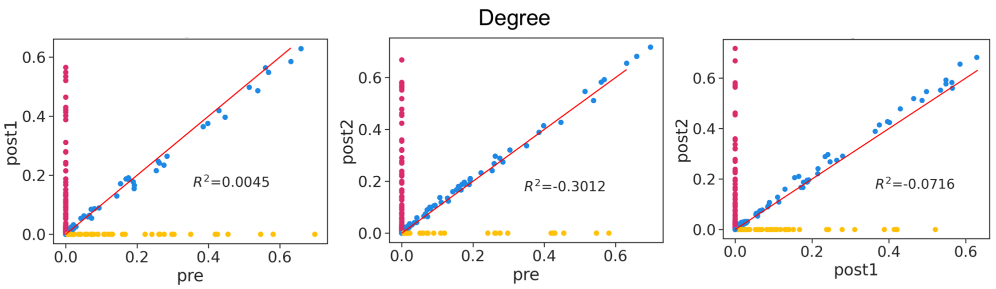


Supplementary Figure 3: Vertex degrees based on ($timepoint_1$, $timepoint_2$) (x- and y-axis, respectively) GRNs inferred from the BC dataset. Coefficient of determination ($R^2$) for $\hat{y} = x$ is reported. Outliers that have degree 0 in one of the GRNs and degree $>$ 100 in another one are highlighted with color. From left to right: (pre,post1); (pre,post2); (post1,post2)

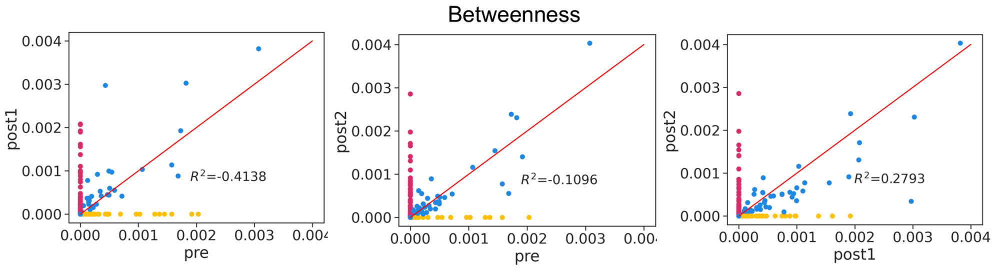


Supplementary Figure 4: Vertex betweenness based on ($timepoint_1$, $timepoint_2$) (x- and y-axis, respectively) GRNs inferred from the BC dataset. Coefficient of determination ($R^2$) for $\hat{y} = x$ is reported. Outliers that have degree 0 in one of the GRNs and degree $>$ 100 in another one are highlighted with color. From left to right: (pre,post1); (pre,post2); (post1,post2)

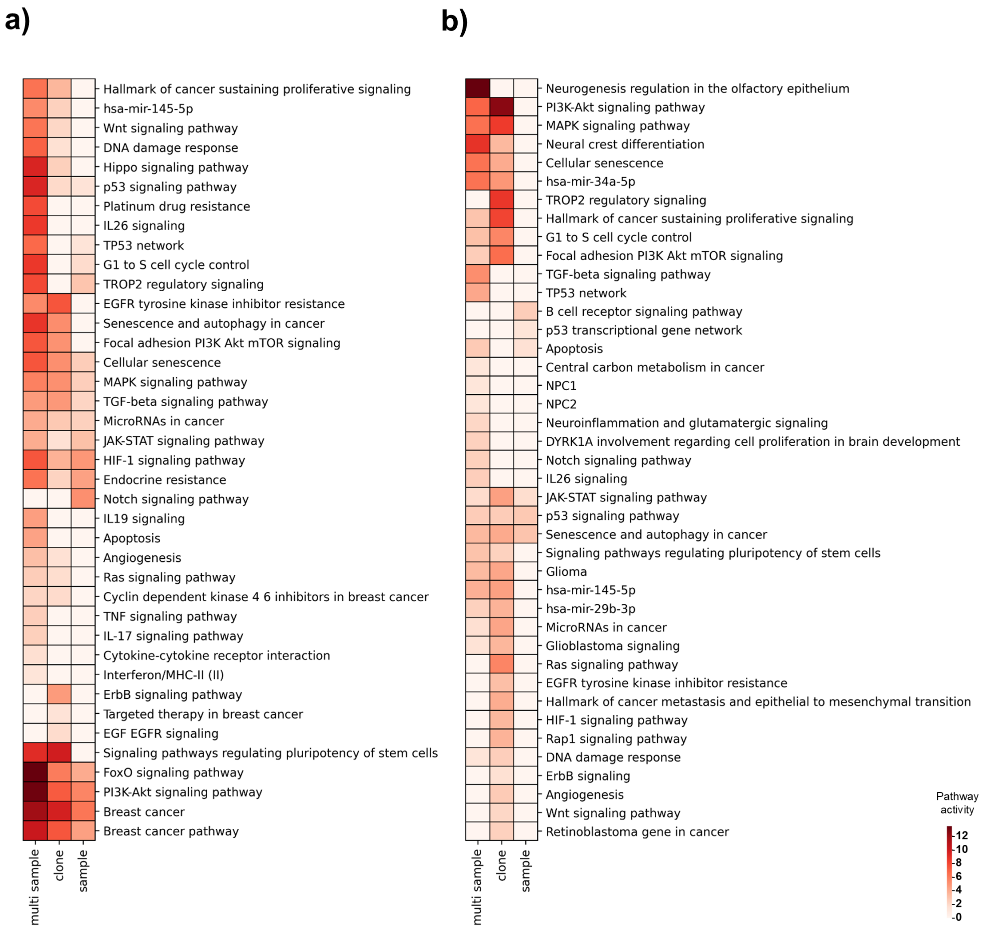


Supplementary Figure 5: (a) Pathways and corresponding pathway enrichment scores specific to each of the three categories in the TNBC analysis. Genesets used for pathway enrichment include the TFs and TGs of every regulon within a specific category (b) Pathways and corresponding pathway enrichment scores specific to each of the three categories in the Glioblastoma analysis. Genesets used for pathway enrichment include the TFs and TGs of every regulon within a specific category

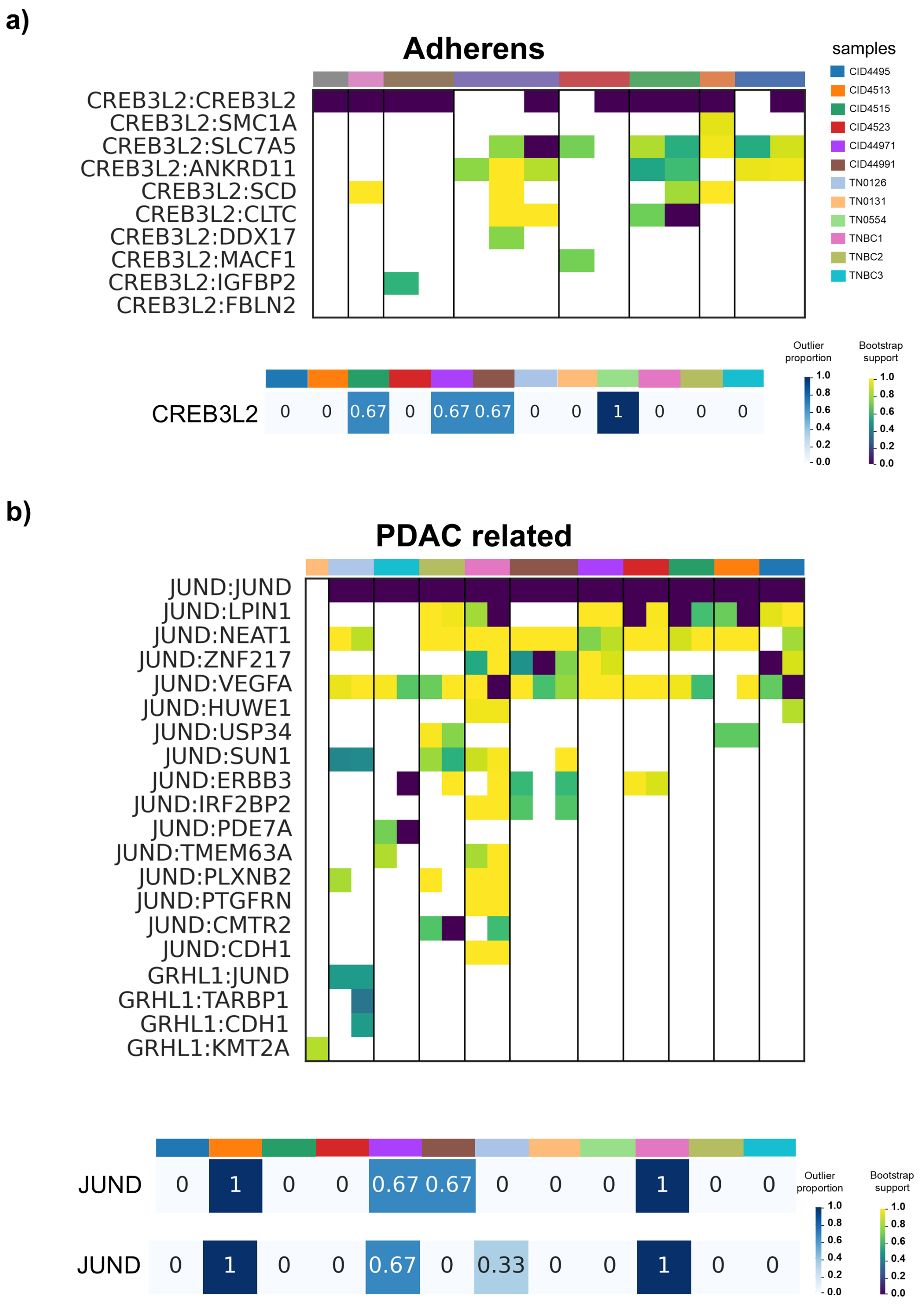
a)
Adherens
samples
CID4495
CID4513
CID4515
CID4523
CID44971
CID44991
TN0126
TN0131
TN0554
TNBC1
TNBC2
TNBC3
CREB3L2:CREB3L2
CREB3L2:SMC1A
CREB3L2:SLC7A5
CREB3L2:ANKRD11
CREB3L2:SCD
CREB3L2:CLTC
CREB3L2:DDX17
CREB3L2:MACF1
CREB3L2:IGFBP2
CREB3L2:FBLN2
CREB3L2 0 0 0.67 0 0.67 0.67 0 0 1 0 0 0
Outlier proportion
Bootstrap support
b)
PDAC related
JUND:JUND
JUND:LPIN1
JUND:NEAT1
JUND:ZNF217
JUND:VEGFA
JUND:HUWE1
JUND:USP34
JUND:SUN1
JUND:ERBB3
JUND:IRF2BP2
JUND:PDE7A
JUND:TMEM63A
JUND:PLXNB2
JUND:PTGFRN
JUND:CMTR2
JUND:CDH1
GRHL1:JUND
GRHL1:TARBP1
GRHL1:CDH1
GRHL1:KMT2A
JUND 0 1 0 0 0.67 0.67 0 0 0 1 0 0
JUND 0 1 0 0 0.67 0 0.33 0 0 1 0 0
Outlier proportion
Bootstrap support

c)

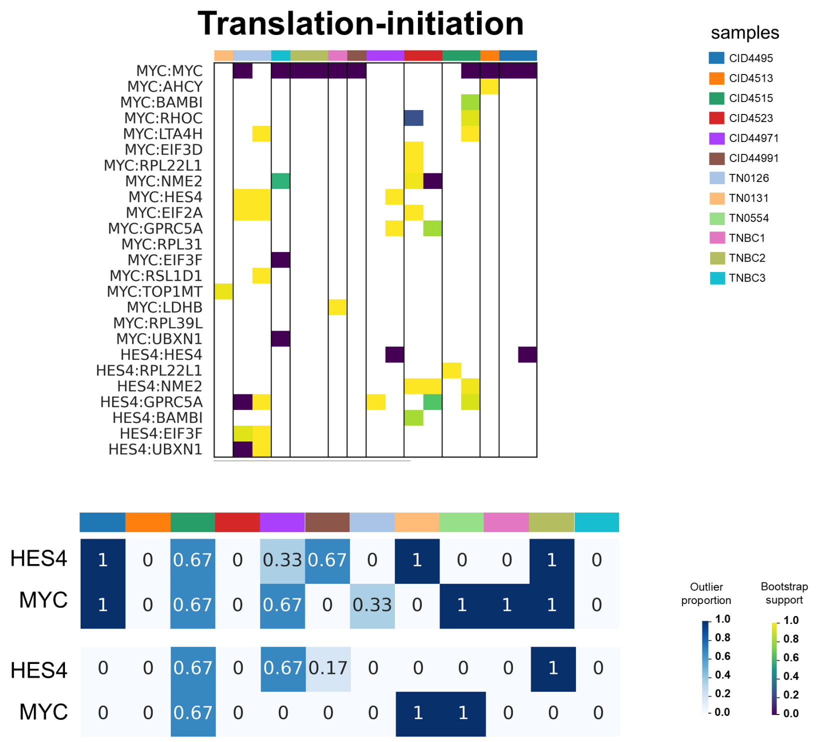


d)

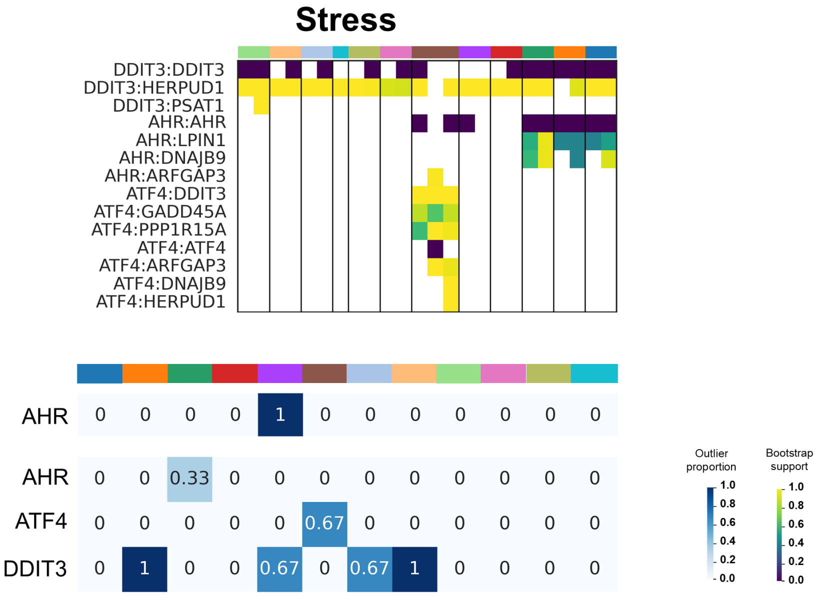


Supplementary Figure 6: (top) Distribution of SCENIC edges across all TNBC samples for meta-programs (a) Adherens (b) PDAC (c) Translation (d) Stress. Each cell represents the corresponding bootstrap value. Sections within each row (sample) represents clones within each sample. (bottom) Distribution of betweenness outliers, across all TNBC samples for meta-programs (a) Adherens (b) PDAC (c) Translation (d) Stress. Each cell represent the fraction of clone pairings a gene has been found to be an outlier in.

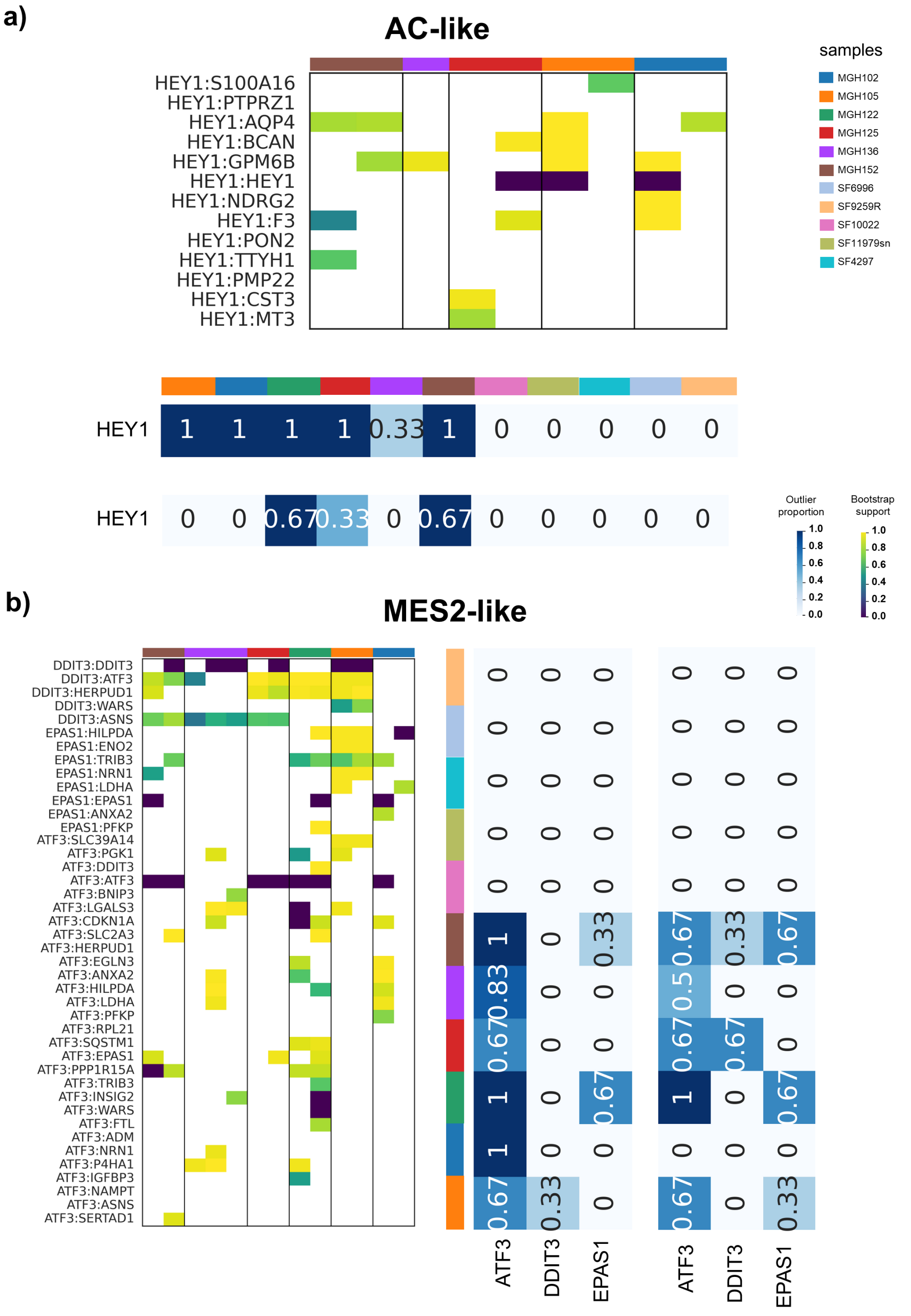


Supplementary Figure 7: (a) (top) Distribution of SCENIC edges across all Glioblastoma samples for meta program AC. Each cell represents the corresponding bootstrap value. Sections within each row (sample) represents clones within each sample. (bottom) Distribution of degree (top) and betweenness (bottom) outliers, across all Glioblastoma samples. Each cell represent the fraction of clone pairings a gene has been found to be an outlier in. (b) (top) Distribution of SCENIC edges across all Glioblastoma samples for meta program MES2. Each cell represents the corresponding bootstrap value. Sections within each row (sample) represents clones within each sample. (bottom) Distribution of degree (top) and betweenness (bottom) outliers, across all Glioblastoma samples. Each cell represent the fraction of clone pairings a gene has been found to be an outlier in.

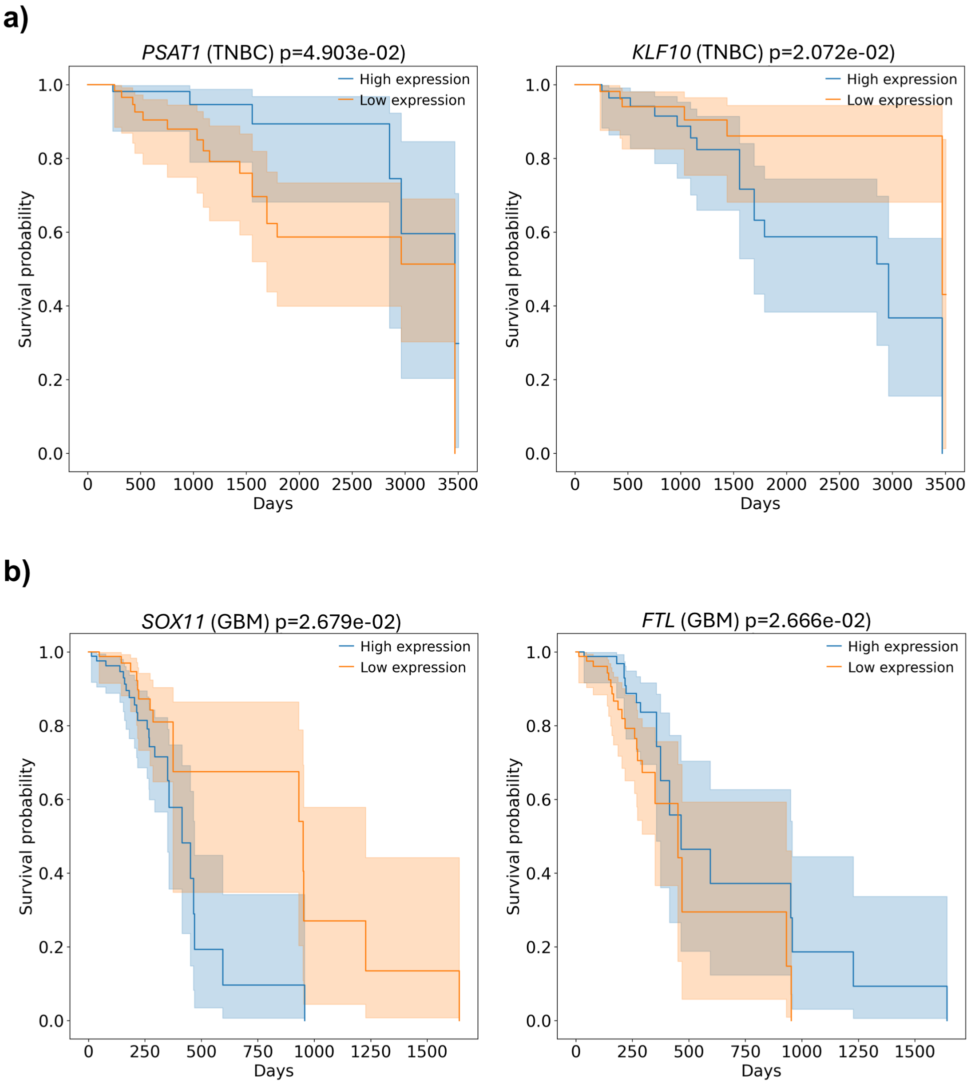


Supplementary Figure 8: (a) Kaplan–Meier survival curves comparing patients stratified by genes *PSAT1* and *KLF10* expression in TNBC patients from TCGA using a median-based threshold across. Patients with expression values greater than or equal to the median were assigned to the high group, and those below the median to the low group. The y-axis represents survival probability, and the x-axis represents time (days). A statistically significant difference in survival between groups was observed (log-rank test, ($p < 0.05$)) (b) Kaplan–Meier survival curves comparing patients stratified by genes *SOX11* and *FTL* expression in GBM patients from TCGA using a median-based threshold across. Patients with expression values greater than or equal to the median were assigned to the high group, and those below the median to the low group. The y-axis represents survival probability, and the x-axis represents time (days). A statistically significant difference in survival between groups was observed (log-rank test, ($p < 0.05$))

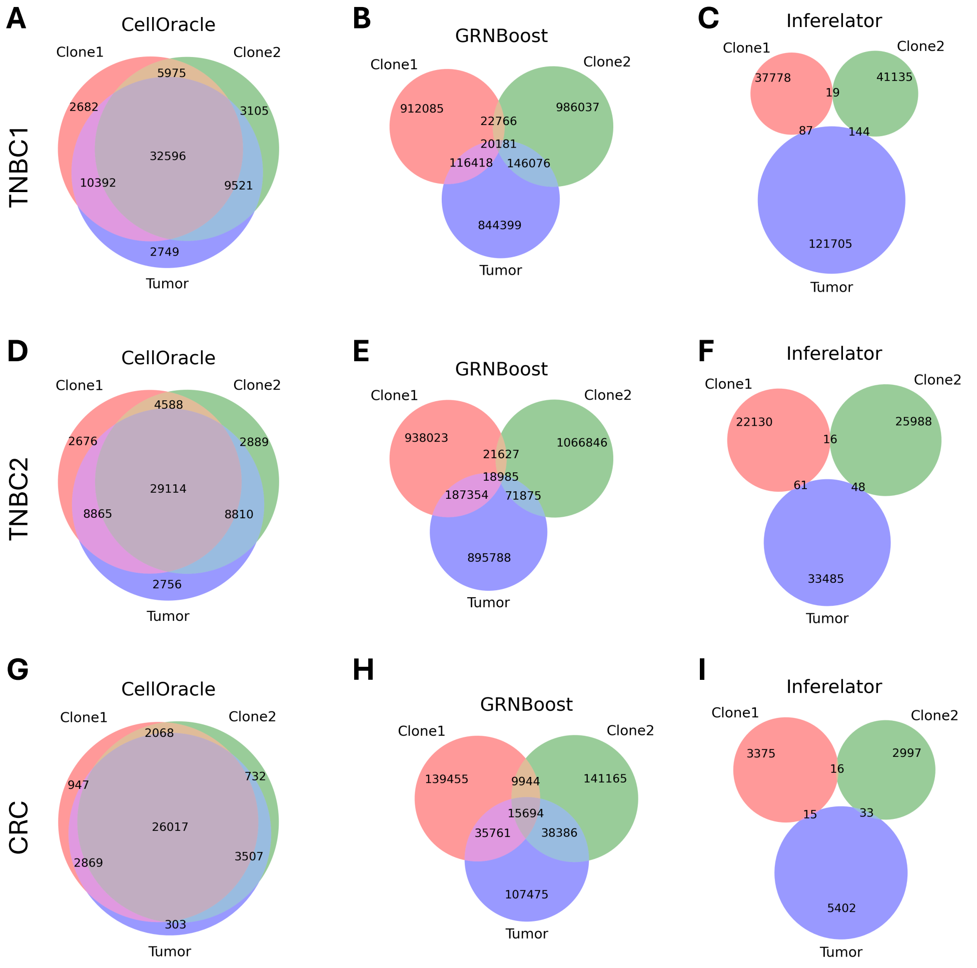


Supplementary Figure 9: Venn diagrams showing unique and shared edges between GRNs inferred from tumor clones 1 and 2, and all tumor cells jointly. Columns correspond to three GRN inference tools: CellOracle **(A, D, G)**, GRNboost2 **(B, E, H)**, and Inferelator **(C, F, I)**. Rows correspond to different samples: TNBC1 **(A-C)**, TNBC2 **(D-F)**, and CRC **(G-I)**
